# Cliffy: robust 16S rRNA classification based on a compressed LCA index

**DOI:** 10.1101/2024.05.25.595899

**Authors:** Omar Ahmed, Christina Boucher, Ben Langmead

**Affiliations:** Johns Hopkins University, Baltimore, MD, USA; University of Florida, Gainesville, FL, USA

**Keywords:** taxonomic, read classification, pangenome, 16S rRNA, document listing

## Abstract

Taxonomic sequence classification is a computational problem central to the study of metagenomics and evolution. Advances in compressed indexing with the *r*-index enable full-text pattern matching against large sequence collections. But the data structures that link pattern sequences to their clades of origin still do not scale well to large collections. Previous work proposed the document array profiles, which use *𝒪* (*rd*) words of space where *r* is the number of maximal-equal letter runs in the Burrows-Wheeler transform and *d* is the number of distinct genomes. The linear dependence on *d* is limiting, since real taxonomies can easily contain 10,000s of leaves or more. We propose a method called cliff compression that reduces this size by a large factor, over 250x when indexing the SILVA 16S rRNA gene database. This method uses Θ(*r* log *d*) words of space in expectation under a random model we propose here. We implemented these ideas in an open source tool called Cliffy that performs efficient taxonomic classification of sequencing reads with respect to a compressed taxonomic index. When applied to simulated 16S rRNA reads, Cliffy’s read-level accuracy is higher than Kraken2’s by 11-18%. Clade abundances are also more accurately predicted by Cliffy compared to Kraken2 and Bracken. Overall, Cliffy is a fast and space-economical extension to compressed full-text indexes, enabling them to perform fast and accurate taxonomic classification queries.

**2012 ACM Subject Classification:** Applied computing *→* Computational genomics

## 1 Introduction

Compressed indexes are increasingly used for read alignment [22, 43, 5] and classification [2, 3] with respect to pangenomes and other collections. These methods exploit the repetitiveness inherent in the Burrows-Wheeler Transform (BWT) of a sequence collection by compressing the BWT into its maximal equal-letter runs [30]. Such approaches can straightforwardly determine if a query string is present or absent in the index [30], but a harder task is to find where the query string occurs. For this, Gagie et al. [20] proposed a subsampling strategy for the suffix array (SA) along with efficient algorithms for “locate” queries, returning all offsets where the substring occurred. This subsampled suffix array fits in a space budget of *𝒪*(*r*) words, where *r* is the number of BWT runs.

Compressed indexes are relevant for taxonomic classification since they enable finding and locating substrings while also compressing away repetitive sequence content. In practice, solving the taxonomic classification problem does not require the full resolution of a locate query. Instead, indexes for taxonomic classification typically support broader queries that either (a) list which genomes the query string occurs in (“document listing”), or (b) find which portion of the taxonomic tree the substring likely originated from (“taxonomic classification”).

We previously introduced a data structure for document listing called the *document array profiles* [1]. This data structure requires 𝒪(*rd*)-space, where *d* is the number of genomes (also called “documents”), and enables document listing in time proportional to the number of documents containing the query substring (*ndoc*). This was the first data structure to enable document listing queries in space proportional to *r* rather than *n*. However, the linear dependence on *d* makes that structure too large for typical taxonomic classification scenarios where *d* can be 10,000 or more. For example, the SILVA NR99 SSU reference database (510,508 sequences, 1.2 GB) spans 9,118 distinct genera, which would require a document array profiles structure larger than 2 TB.

Here, we develop new methods that drastically reduce the space usage. In so doing, we sacrifice the ability to perform full document listing, but we retain the ability to perform accurate taxonomic classification. Detecting and measuring alleles of the 16S rRNA gene allows for the study of bacterial communities, a common task since its introduction in the 1970s [50]. The gene contains both conserved and hypervariable regions, allowing for primer design (in conserved portions) [15, 48] and measurement of evolutionary distance (in variable portions) [25, 45]. Projects like the Human Microbiome Project [11], Earth Microbiome Project [47] and MetaSub [46] have used 16S sequencing to characterize the microbes present in the gut, skin, soil, urban environments, and water. In clinical settings, the 16S rRNA gene has been used, for example, to study spatial patterns of microbial infection in lungs of cystic fibrosis patients [49] as well as microbial communities in the saliva of Behçet’s syndrome patients [10].

Existing software tools for 16S rRNA classification use databases such as Greengenes [13], RDP [29], and SILVA [40] as the reference collection. In 2018, Almeida et al. [4] performed a benchmarking study that showed the QIIME2 [6] was more accurate than other tools for quantifying reads at the genus and family levels, but was computationally very expensive, with time and memory usage 2 and 30 times higher respectively than other tools [4]. More recently, Lu et al. [28] extended the popular shotgun metagenomic classifier Kraken2 [51] to support 16S rRNA databases, showing that their method was up to 300 times faster than QIIME2 while achieving higher accuracy and being the only tool to provide per-read classifications.

Our methods are implemented in a new tool called *Cliffy* that performs per-read 16S rRNA gene classifications using a compressed full-text index. Cliffy uses a novel compression scheme to reduce the size of the document array profiles [1] by two orders of magnitude to make it practical for 16S rRNA classification. In addition, Cliffy incorporates strategies such minimizer digestion and a backward search lookup table to speed up its queries. Using a similar approach to the benchmarking study by Almeida et al. [4], we generate realistic read datasets from different biomes and show that Cliffy consistently outperforms Kraken2 in terms of per-read accuracy as well as genus abundance profiling.

## 2 Methods

### Preliminaries

We define a string *T* [1..*n*] with length *n* as a concatenation of characters *T* [1] *…·T* [*n*] drawn from alphabet Σ of size *σ*. The *empty* string, denoted *ε*, is the only string with length 0. We assume *T* ends with $ ∉ Σ, a special symbol lexicographically smaller than all the symbols in Σ.

Given two integers 1*≤i ≤j ≤n*, we denote *T* [*i*..*j*] = *T* [*i*] … *T* [*j*] the *substring* of *T* spanning positions *i* to *j*. Otherwise if *i > j*, we define *T* [*i*..*j*] = *ε*. We refer to *T* [*i*..*n*] as the *i*-th *suffix* of *T* and *T* [1..*i*] as the *i*-th *prefix* of *T* . Given two strings *T* [1..*n*] and *S*[1..*m*], lcp(*S, T*) denotes the length of the longest common prefix (LCP) of *T* and *S*.

### Suffix array, inverse suffix array, and longest common prefix array

Given a string *T*, the *suffix array* [31] SA_*T*_ [1..*n*] is the permutation of {1, …, *n*} that lexicographically sorts the suffixes of *T* . The *inverse suffix array* ISA_*T*_ [1..*n*] is the inverse permutation of SA_*T*_ [1..*n*], i.e., for all *i* = 1, …, *n* ISA_*T*_ [SA_*T*_ [*i*]] = *i*. The *longest common prefix* array LCP_*T*_ [1..*n*] stores the length of the LCP between lexicographically consecutive suffixes of *T*, formally, LCP[1] = 0 and for all *i* = 2, …, *n* LCP[*i*] = lcp(*T* [SA_*T*_ [*i −* 1]..*n*], *T* [SA_*S*_[*i*]..*n*]).

### Burrows-Wheeler transform

Given a string *T*, the *Burrows-Wheeler transform* [8] BWT_*T*_ [1..*n*] is a reversible permutation of *T* defined as the last column of the matrix of the lexicographically sorted rotations of *T* . When *T* is terminated by $ we can define for all *i* = 1, …, *n*, BWT_*T*_ [*i*] = *T* [SA_*T*_ [*i*] *−* 1] where *T* [0] = *T* [*n*]. LF*-mapping* is the the permutation of 1, …, *n* that maps every character in the BWT_*T*_ to its predecessor in text order, formally LF[*i*] = ISA_*T*_ [SA_*T*_ [*i*] *−* 1] for *i* where SA_*T*_ ≠ 1 and LF[*i*] = ISA_*T*_ [*n*] when SA_*T*_ = 1. We define *r* as the number of maximal equal-letter runs of BWT_*T*_ . When clear from the context we refer to SA_*T*_, ISA_*T*_, LCP_*T*_, and BWT_*T*_ as SA, ISA, LCP, and BWT respectively.

### *r*-index

The *r*-index [21] is a full-text index based on the run-length encoded BWT and the SA entries sampled at run boundaries. Given a text *T* [1..*n*] and a pattern *P* [1..*m*] the *r*-index can locate all occurrences of *P* in *S* in *𝒪* (*m* log log_*w*_(*σ* + *n/r*) + *occs* log log_*w*_(*n/r*)) time and *𝒪* (*r*) words of space, where *occs* is the number of occurrences of *P* in *S*.

### Document array and document array profiles

We denote a collection of documents (i.e. strings) with *𝒟* = {*T*_1_, …, *T*_*d*_} and we denote a concatenation of documents with *𝒯* [1..*n*] = *T*_1_ *· · · T*_*d*_. The *document array* [34] DA[1..*n*] stores for each position *i* the document index of *𝒯* [SA_*𝒯*_ [*i*]..*n*]. The document array profiles [1] is a data structure that enables the *r*-index to perform *document listing* queries.

#### ▶ Problem 1.

*Document listing: Given a collection* 𝒟 = {*T*_1_, …, *T*_*d*_} *and a pattern P, return the set of documents ℒ ⊆ 𝒟 where P occurs.*

Given an *r*-index for the concatenated text *𝒯* [1..*n*], the document array profiles supports document listing for a pattern *P* [1..*m*] in *𝒪* (*m* log log_*w*_(*σ* +*n/r*)+*ndoc*) time and *𝒪* (*rd*)-space, where *ndoc* is the number of documents containing the pattern *P* .

#### ▶ Definition 1.

*(From [1]) Document array profiles: For all positions* 1*≤*LF(*i*) *≤n where i is a run head or tail in the* BWT*of T*, *the document array profile P*_*DA*_[*i*][1..*d*] *stores for each position j* = 1, …, *d the length of the longest common prefix between 𝒯* [*SA*[*i*]..*n*] *and all suffixes of document T*_*j*_.

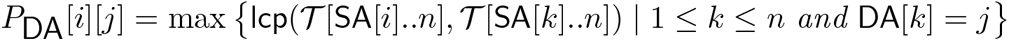

### Document listing query using document array profiles

In our previous work [1], we demonstrated how to compute a document listing for a pattern *P* using the document array profiles. The algorithm requires that we store *P*_*DA*_[*LF* (*i*)] for all *i* corresponding to a BWT run head or tail, following the “toehold lemma” strategy [39], i.e., the stored rows are sufficient to recover the document listing for a query pattern *P* .

Given a query pattern *P*, we perform backward search for *P* and store a pointer to a “sampled” row *P*_*DA*_[*i*] of the document array profiles where BWT[*i*] = *P* [*j*] where *j* is the last character of *P* that was searched. At each step of the backward search, we either sample a new row or simply increment a counter variable *k* by 1. If the backward search range BWT[*s*..*e*] spans a BWT run head or tail, we sample a new row *P*_*DA*_[*i*] where *s≤i < e* and reset *k* to 0. Otherwise, if the backward search range is contained within a BWT run, we simply increment *k* by 1.

In order to generate the document listing, we add *k* to each value in the “sampled” row *P*_*DA*_[*i*][1..*d*] and return all values of *j* where *P*_*DA*_[*i*][*j*] *≥*|*P*|. Assuming the LCP values in *P*_*DA*_[*i*][1..*d*] are stored in sorted order, we only need to scan the first *ndoc* + 1 values to generate the document listing. As previously mentioned, the document array profiles supports document listing for a pattern *P* [1..*m*] in *𝒪* (*m* log log_*w*_(*σ* + *n/r*) + *ndoc*) time and *𝒪* (*rd*)-space. More details on the proof for these bounds can be found in [1].

### Taxonomic classification with the pattern lowest common ancestor query

We aim to classify a sequence according to where it likely originated from in the tree of life, represented by a *taxonomy*. Let a taxonomy *R* be a rooted multiway tree with *d* leaves. Given a taxonomy *R*, we define the collection of documents of the taxonomy, denoted by {*R*_1_, …, *R*_*d*_} as the collection of reference sequences of the taxonomic clades at the leaves of the taxonomy (Figure 1a). Leaves of the taxonomy *R* map in a one-to-one fashion with the documents. Internal nodes represent common ancestors, and do not have directly associated strings; rather, they are implicitly associated with the union of the documents in the leaves in their subtree. Since a one-to-one mapping exists between leaves and documents, we often refer to these interchangeably.

**Figure 1.**
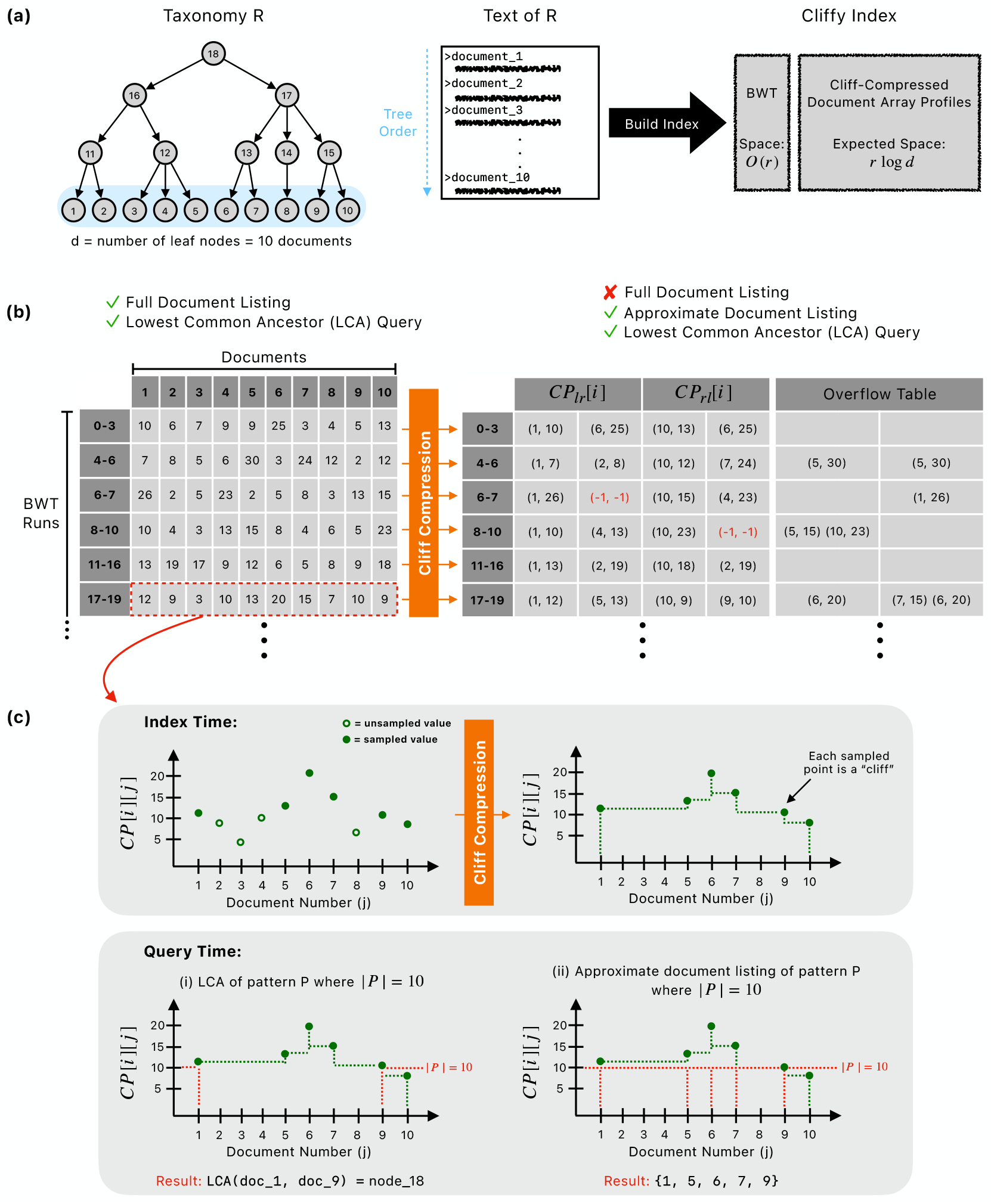
(a) Shows an example taxonomy where *d* = 10 and an example tree order that guides the generation of the text of the taxonomy prior to building an index. (b) Visualizes the difference between the document array profiles and the cliff-compressed document array profiles. (c) Using a profile (i.e. row of document array profiles) as an example, visualizes cliff-compression at the index building step, followed by how each profile is used at query time for either the lowest common ancestor query or approximate document listing.

#### ▶ Definition 2.

*The occurrences of a string P in taxonomy R is the set of leaves ℱ where P occurs as a substring of the associated document*.

We summarize where a string *P* originated using a *pattern lowest common ancestor* query:

#### ▶ Problem 2.

*Pattern lowest common ancestor (PLCA): Given a taxonomy R and string P*, *return the deepest node X such that all of P’s occurrences are in leaves of X’s subtree*.

In order to process a set of occurrences of a pattern *P* in a taxonomy, we define the *lowest common ancestor* query:

#### ▶ Definition 3.

*Lowest common ancestor (LCA): Given a taxonomy R and set of nodes ℱ from R, the LCA of ℱ is the deepest node X in R where all nodes in ℱ are present in X’s subtree*.

PLCA can be solved by first identifying all occurrences of a pattern *P* in the leaves, then computing the LCA of those leaves. However, a simpler approach becomes possible if we first consider the leaf nodes to be in a particular order [19]:

#### ▶ Definition 4.

*Tree order: A tree order of the d leaves of taxonomy R is any ordering such that all leaves descended from any internal node are consecutive in the order*.

Alternately, we could say that the tree order ensures that descendants of sibling nodes in the tree cannot be interleaved in the order. From here on, we assume that the concatenated text *𝒯* [1..*n*] respects tree order for the taxonomy *R*. We also assume that the document ids associated with the leaves follow this order, i.e., the leftmost (earliest) leaf is document 1, the next is document 2, etc.

#### ▶ Lemma 5.

*(From [19]) Assuming tree order, the PLCA query can be answered by computing the LCA of only the leftmost (earliest) and rightmost (latest) leaf nodes containing P in the order*.

**Proof**. The proof is by contradiction. We assume a pattern *P* has occurrences ℱ= {ℱ_1_, …, ℱ_*n*_ } in taxonomy *R*. Further, we assume the occurrences are sorted by document ids, such that ℱ_1_ is an occurrence in the leftmost of the documents that *P* occurs in and ℱ_*n*_ is an occurrence in the rightmost document that *P* occurs in. We let LCA() denote the LCA of the full set of occurrences, and LCA(ℱ_1_, ℱ_*n*_) denote the LCA of the leftmost and rightmost occurrences.

We presume LCA(ℱ) ≠ LCA(ℱ_1_, ℱ_*n*_). Then it follows that there is a node 𝒳 ⊆ ℱ\{ ℱ_1_, ℱ_*n*_} that occurs either in a document prior to the one matched in occurrence ℱ_1_, or in a document after the one matched in occurrence ℱ_*n*_. This implies the ordering allows for interleaving of the descendants of sibling nodes, contradicting the assumption of tree ordering. ▶

#### ▶ Lemma 6.

*Given an r-index for a tree-ordered text of a taxonomy 𝒯* (*R*) = *R*_1_ *R*_*d*_ *extended with the document array profiles P*_*DA*_, *we can compute the PLCA of the pattern P* [1..*m*] *in 𝒪* (*m* log log_*w*_(*σ* + *n/r*) + *ndoc*)*-time and 𝒪* (*rd*)*-words of space*.

**Proof**. As summarized in the preliminaries, we previously showed how to compute the document listing for any length substring in *𝒪* (*m* log log_*w*_(*σ* +*n/r*)+*ndoc*)-time by combining backward search with a scan of a “sampled” row of the document array profiles [1]. Since we have a tree order, the PLCA for the pattern *P* can be computed as the LCA of the leftmost and rightmost leaves for which *P* occurs in their associated document. The LCA of these two leaves is then computed with an auxiliary LCA data structure requiring 2*d* + *o*(*d*) bits of space with *𝒪* (1) query time [18, 35]. These additional operations are both constant time, *𝒪* (1), therefore, total time complexity is *𝒪* (*m* log log_*w*_(*σ* + *n/r*) + *d*). ▶

### Shrinking the document array profiles with cliff compression

In the previous section, we proved that we can compute the PLCA for a given pattern using the document array profiles, however, it requires *𝒪* (*rd*)-space. We note that in practical settings, *d* can reach tens of thousands of species (documents). We propose a strategy that we refer to as *cliff compression*, which has an average space usage of Θ(*r* log *d*) while still allowing for the exact computation of PLCAs. The insight is that to determine the leftmost document containing an *𝓁*-length match, it is sufficient to identify the smallest *j* such that *P*_*DA*_[*i*][*j*] *≥𝓁*, where *i* represents the current BWT offset. Similarly, when we ask which is the rightmost document containing an *𝓁*-length match, it is sufficient to know the largest *j* such that *P*_*DA*_[*i*][*j*] *≥ 𝓁*. As a result, we only need to store the maxima-so-far, i.e., max(*P*_*DA*_[*i*][1..*j*]) and max(*P*_*DA*_[*i*][*j*..*d*]) for all *j* when they first occur. Though we must store these for both the left-to-right and right-to-left maxima-so-far, the number of instances *m* where the maxima-so-far changes and we need to store an additional value is small in practice (*m≪ d*), making them significantly smaller to store compared to the full document array profile row *P*_*DA*_[*i*][1..*d*].

For added intuition, we visualize the profile *P*_*DA*_[*i*][1..*d*] (Figure 1c) with the maxima-so-far step functions drawn as dotted lines. The leftmost and rightmost documents containing a pattern *P* corresponds to the *j* values where a height- |*P*| horizontal line crosses the maxima-so-far function. The LCA of the documents at the crossings is the PLCA. Longer matches correspond to higher horizontal lines, which tend to cross a narrower portion of the maxima-so-far function and so leads to deeper PLCAs.

#### ▶ Definition 7.

*Cliff compression: Given the full document array profiles P*_*DA*_, *we cliff-compress each row by first scanning P*_*DA*_[*i*][1..*d*] *from left to right and storing all* (*j, P*_*DA*_[*i*][*j*]) *pairs where j* = 1 *or P*_*DA*_[*i*][*j*] *>* max (*P*_*DA*_[*i*][1..*j−*1]). *We store these in an array denoted as CP*_*𝓁r*_[*i*], *in ascending order by j. We also scan P*_*DA*_[*i*][1..*d*] *from right to left and store all* (*j, P*_*DA*_[*i*][*j*]) *pairs, where j* = *d or P*_*DA*_[*i*][*j*] *>* max (*P*_*DA*_[*i*][*j* + 1…*d*]). *We collect these in an array called CP*_*r𝓁*_[*i*], *in descending order by j. CP*_*𝓁r*_ *and CP*_*r𝓁*_ *together constitute the cliff-compressed document array profiles*.

We can compute PLCAs using only the cliff-compressed document array profiles.

#### ▶ Theorem 8.

*Given the r-index built over the tree-ordered text of a taxonomy 𝒯* (*R*) = *R*_1_ *R*_*d*_, *extended with the cliff-compressed document array profiles CP*_*𝓁r*_ *and CP*_*r𝓁*_, *we can compute the PLCA for a pattern P in𝒪* (*m* log log_*w*_(*σ* + *n/r*) + *d*) *time in 𝒪* (*rd*) *words of space*.

**Proof**. We adapt the query algorithm given in the preliminaries to work with cliff-compressed document array profiles. Instead of storing *P*_*DA*_[*LF* [*i*]] at each head or tail, we store the pair (*CP*_*𝓁r*_[*LF* [*i*]], *CP*_*r𝓁*_[*LF* [*i*]]). The space usage remains *𝒪* (*rd*) in the worst case. During backward search, when the range includes a run head or tail, we retrieve (*CP*_*𝓁r*_[*LF* [*i*]], *CP*_*r𝓁*_[*LF* [*i*]]) for *i* corresponding to the (contained) head or tail. When the range falls entirely within a run, we obtain our new sample by incrementing each *P*_*DA*_[*i*][·] value (i.e. LCP) stored in the *CP*_*r𝓁*_[*i*] and *CP*_*𝓁r*_[*i*] lists from the previous step by 1.

After backward search, we find the leftmost and rightmost documents containing *P* . To find the leftmost, we scan the pairs of (*j, P*_*DA*_[*i*][*j*]) ∈ *CP*_*𝓁r*_[*i*], retrieving the first value *j* where *P*_*DA*_[*i*][*j*] *≥* |*P* |. By definition of *CP*_*𝓁r*_, this is the leftmost document containing *P* . Next, we scan the pairs (*j, P*_*DA*_[*i*][*j*]) *∈ CP*_*r𝓁*_[*i*] and retrieve the first value *j* such that *P*_*DA*_[*i*][*j*] *≥* |*P* | (recalling that elements of *CP*_*r𝓁*_[*i*] are in descending order by *j*). By definition of *CP*_*r𝓁*_, this is the rightmost document containing *P* . These scans each require *𝒪* (*d*)-time.

Finally, we query the LCA data structure with leftmost and rightmost documents in *𝒪* (1)-time [18, 35] to complete the PLCA query. Thus, we can compute the PLCA for a pattern *P* using the cliff compressed version of *P*_*DA*_ in *𝒪* (*m* log log_*w*_(*σ* + *n/r*) + *d*)-time. ▶

### Random model for cliff compression ratio

Cliff compression does not improve worst-case space usage compared to the original document array profiles—both require *𝒪* (*rd*) words of space, with the worst-case for cliff compression of *CP*_*𝓁r*_[*i*] (resp. *CP*_*r𝓁*_[*i*]) being the case where the LCP values in *P*_*DA*_[*i*] are strictly increasing (resp. decreasing). However, we now show that the average-case space complexity when applying cliff compression to the document array profiles is significantly smaller. To accomplish this, we begin by proposing a random model that leads to the *average* compression ratio of cliff compression, and then we will describe the links between the random model and the cliff-compression scenario.

We model the values in a row *i* of the document array profiles (*P*_*DA*_[*i*]) as a random permutation of the *d* integers 0, 1, …, *d−* 1. We then consider the “minimum-so-far” computed over the permutation, by computing the minimum over all non-empty prefixes of the permutation. We model the number of values we must store in the *CP*_*𝓁r*_[*i*] or *CP*_*r𝓁*_[*i*] list as the number of times that the the minimum-so-far sequence decreases, plus one. Adding one accounts for the fact that we must always store at least one pair in the array. Note that once the minimum-so-far reaches 0 it cannot decrease further.

In practice, the document array profiles deviate from this model. Most importantly: (1) the documents in the taxonomy are related to each other, violating any assumptions of uniform probability of different LCP values, and (2) the values in *P*_*DA*_[*i*] are not all distinct as in the simple model. Nonetheless, we found that the model was effective at predicting the number of pairs in the cliff-compressed document array profiles (Table 1).

**Table 1.** Size of the Cliffy indexes for the SILVA SSU NR99 (version 138.1) database which contains 510, 508 rRNA sequences. Each distinct genera in the SILVA database is treated as a distinct document and there are 9,118 genera (*d* = 9118). The table also lists for each digestion approach the average number of pairs stored for each cliff-compressed profile. The expected number pairs as discussed in the Methods section is based on the Harmonic series sum, *H*_9118_ + 1 = 10.695.

| Reference Digestion | Index Size |  | Reduction Ratio | Avg # of Pairs |
| --- | --- | --- | --- | --- |
|  | Document Array Profiles<br>(Ahmed et al., 2023) | Cliffy<br>(this paper) |  |  |
| No Digestion | 2265 GB | 9.014 GB | 251X | 7.095 |
| DNA minimizer | 1955 GB | 7.858 GB | 249X | 7.211 |
| Minimizer | 1172 GB | 4.266 GB | 275X | 5.137 |

Given a random permutation of the *n* integers 0, 1, …, *n−* 1, we define the random variable *S*_*n*_ as the number of minima-so-far decreases before reaching 0. The base case is *S*_0_ = 0, i.e., if the leftmost value in the permutation is 0 then the minimum-so-far is uniformly 0 with no decreases. Another easy case is *S*_1_ = 1, i.e., if the leftmost value is 1 then exactly one decrease (to 0) will occur. For *S*_2_, we must consider two cases: we reach 1 before reaching 0, or we reach 0 before reaching 1. In the first case, *S*_2_ = *S*_1_ + 1 = 2. In the second, *S*_2_ = *S*_0_ + 1 = 1. The cases are equally likely since the permutation is random. Therefore, we have **E**[*S*_2_] = 1 + 1*/*2 *·* (**E**[*S*_1_] + **E**[*S*_0_]) = 3*/*2. In general, we have:

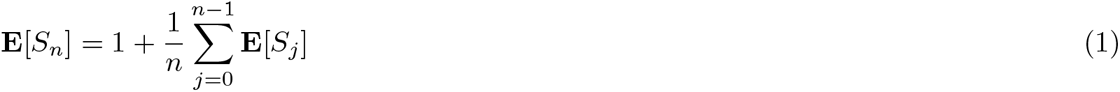

A closed form of the above formula equals the *n*^th^ harmonic number:

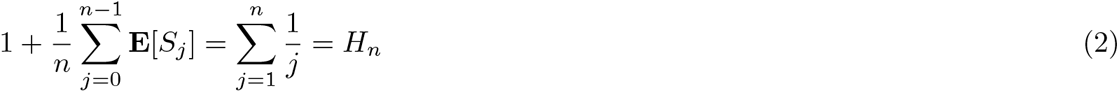

We prove this formula using induction in Appendix A.1. Finally, we note that *H*_*n*_ grows with Θ(log *n*), as is proved in Appendix A.2. That is, the expected number of decreases in the minima-so-far of a random permutation the integers in 0, 1, …, *n−* 1 grows with log *n*. We summarize this finding in the following theorem.

#### ▶ Theorem 9.

*Given the r-index built over the tree-ordered text of a taxonomy 𝒯* (*R*) = *R*_1_ *R*_*d*_, *extended with cliff-compressed document array profiles where the LCPs follow the distribution of random permutations described in the above model, we can compute the PLCA for pattern P in 𝒪* (*m* log log_*w*_(*σ* + *n/r*) + *d*) *time and* Θ(*r* log *d*) *words of space*.

**Proof**. In theorem 8, we showed that we can compute the PLCA for a pattern *P* using the cliff-compressed document array profiles in *𝒪* (*m* log log_*w*_(*σ* + *n/r*) + *d*)-time in *𝒪* (*rd*)-space. Assuming the LCP values in each row of the original document array profiles follow the distribution of random permutations, we prove there is a lower average space bound. We showed in Appendix A.2 that the expected number of pairs that will be stored in *CP*_*𝓁r*_[*i*] and *CP*_*r𝓁*_[*i*] is Θ(log *d*), therefore, the total average space for the cliff-compressed document array profiles is Θ(*r* log *d*). ▶

### Approximate document listing

The cliff-compressed document array profiles enables us to perform an additional type of query called *approximate document listing*. Rather than returning the PLCA of the pattern *P*, this query returns a subset of the documents in the true document listing for *P* .

#### ▶ Definition 10.

*Approximate document listing: Given a sampled cliff-compressed profile, CP*_*𝓁r*_[*i*] *and CP*_*r𝓁*_[*i*], *and pattern P*, *the approximate document listing of P is obtained by scanning all the pairs* (*j, P*_*DA*_[*i*][*j*]) *in both CP*_*𝓁r*_[*i*] *and CP*_*r𝓁*_[*i*]_*R*_ *and returning the set of j where CP*_*lr*,*rl*_[*i*][*j*] *≥* |*P* |.

The motivation for this query is to avoid the potential for vague PLCAs, especially those near the root of the taxonomy. For example, suppose we have a substring *M* from a read that occurs in *n* documents, the PLCA of *M* would be the root of taxonomy even if *n−* 1 of the occurrences of *M* were at the left-end (earliest) of the tree-order and the last occurrence was in the rightmost (latest) document in the tree-order. This illustrates a scenario when the PLCA of *M* gives an unclear classification even though *M* most likely originated from a document in the left side (earlier) of the tree-order.

We note that both the PLCA and approximate document listing queries can be performed by using the same cliff-compressed profile. Despite the fact that it fails to be a full document listing, we show in Results that the approximate document listing can be a more informative query for classification compared to the PLCA in practice.

### Classification methods in Cliffy

Cliffy can perform both read-level classification and abundance profiling. Given a sequencing read, Cliffy begins at the end of the read and performs maximal left-extension using backward search on the tree-ordered text of the taxonomy. Once the backward search range is empty, Cliffy resets the range to the full BWT at position *i*, where the previous maximal left-extension ended. Thus, the features for classification used by Cliffy are the exact matches (of varying lengths) between the read and the reference database. Each exact match will have an associated cliff-compressed profile that allows us to either compute the PLCA or approximate document listing. For example, if we would like to classify at the genus level of the taxonomy and there are *d* distinct genera, then we initialize a vote array *V* [1..*d*] with all zeroes. Next, we iterate through each exact match *M*, and compute the PLCA for *M* that returns the leftmost *𝓁* and rightmost *r* occurrence of *M* in the taxonomy. We add 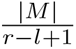 to all entries between *V*[*l*] and *V* [*r*], inclusive on both ends, in order to distribute the length of the exact match among all of the leaf nodes in the subtree of the PLCA of *M* . If we would like to use approximate document listing instead to classify the read, then we consider each exact match *M* and retrieve the approximate document listing *ℒ* and then add 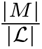 to the entries in *V* corresponding to documents *j*, where *j* ∈ ℒ.

Finally, the read is classified by identifying the document *j* with the maximum value *V* [*j*] in *V* . In order to perform abundance profiling at the genera level, we classify each read individually and return the proportion of reads classified to each genus.

### Running Kraken2/Bracken for classification and profiling

In our experiments, we compared against Kraken2 for read-level classification and the Kraken2+Braken pipeline for abundance profiling. The commands to build the Kraken2/Bracken indexes as well as using them for classification are listed in the Appendix A.3.

### Simulating realistic 16S rRNA datasets

In order to benchmark Cliffy and Kraken2, we simulated 16S rRNA reads using an approach of Almeida et al. [4]. More specifically, we implemented and used a Python tool called `MicrobeMixer` that allows us to simulate realistic 16S rRNA reads for different biomes. The commands used can be found in Appendix A.4.

Similar to Almeida et al. [4], we used EBI’s MGnify API [41] to query public metagenomic datasets from three different environments (Human Gut, Aquatic and Soil) to identify the top 100 most common genera of bacteria. Next, using the SILVA SSU NR99 database, we take all of the reference sequences for the top 100 genera and extract the sequences in different hypervariable regions (V1-V2, V3-V4, V4, V4-V5) (exact primers can be found in Appendix A.4) using in-silico PCR where we allow up to 3 mismatches in the primer sequence. Next, paired-end sequencing reads 250 bp in length are simulated from these extracted regions using *ART* [23]. Finally, `MicrobeMixer` sampled reads from each of the top 100 genera, mimicking their abundance in the public data, to generate a dataset containing 10 million paired-end reads.

### Reference digestion with minimizers

Cliffy uses minimizer digestion similar to SPUMONI 2 [3] in order to reduce of the size of text prior to indexing. The first type of digestion is the classical case and we refer to it as “DNA minimizer”, where we have a small (*k*) and large (*w*) window size. The *w− k* + 1 *k*-mers in large window are hashed and the minimizer is the *k*-mer with the smallest hash value. As Cliffy digests the text, it only concatenates minimizers that are distinct from the preceding one. The second type of digestion treats minimizers themselves as characters opposed to using a “DNA minimizer” which is a concatenation of *k* characters. This strategy was previously implemented in assembly-based tools such as mdbg [16] and ntJoin [12]. Cliffy uses 4 for the value of *k* since with four DNA nucleotides (A, C, G, T) there are 256 possible 4-mers which can all be represented with 1 byte. Both minimizer approaches allow Cliffy to reduce the size of the text prior to indexing which in turn reduces the size of the index.

### Accelerated querying optimizations in Cliffy

There are two main optimizations in Cliffy that accelerate the query: ftab and minimizer digestion. Firstly, ftab is a lookup table used in backward search to bypass a pre-specified amount of backward search steps inspired by Bowtie2 [27] and Rowbowt [33]. In Cliffy, the ftab the stores the backward search ranges (start and end positions) for all possible 3-mers in the minimizer alphabet. Then at query-time, it begins by checking if the 3-mer at the end of the read is present in the text using the ftab, if so, it moves to the 4th character and proceeds with backward search as normal. Whenever Cliffy resets the backward search range to the full BWT, it will check to see if the ftab can be used to accelerate the query.

Secondly, as previously discussed, Cliffy uses minimizer digestion to reduce both the input text and the reads to shorter sequences. Importantly, this digestion accelerates the query since Cliffy treats each minimizer as its own character; thereby, if *k* = 4, Cliffy can replace four backward search steps with one.

## 3 Results

We performed the experiments on an Intel Xeon gold 6248R 24-core 3GHz processor with 1,500 GB of RAM. We ran on a 64-bit Linux platform. Time was measured using GNU `time`. Source code and the experimental scripts using Snakemake [26] are available on GitHub (see Software availability).

### Datasets

We simulated 12 realistic read datasets from three different environments (“biomes”) and four different hypervariable regions within the 16s rRNA gene. Each read set consisted of 10 million 250 bp Illumina paired-end reads. The exact simulation parameters can found in Appendix A.4.

### Comparable Methods

Other taxonomic classification methods can be broadly categorized as: (1) *k*-mer based approaches that index the *k*-mers from the reference, extract *k*-mers from the read, and use these as keys for finding longer matches [51, 38, 7, 42, 6], and (2) full-text approaches that create a full-text index to query with substrings of any length [24, 44, 3, 32].

A *k*-mer based index typically maps *k*-mers to a summary indicating which taxonomic clades contain the *k*-mer. Popular tools like Kraken [52] and Kraken2 [51] store the lowest common ancestor of the genomes that include the *k*-mer (or minimizer of the *k*-mer, in the case of Kraken2). In contrast, full-text based approaches [24, 44, 3], including Cliffy, maintain data structures over the entire reference database to rapidly perform document listing queries on substrings from the sequencing read.

We compare here to Kraken2 since it is the only other method designed for classifying individual 16S rRNA reads rather than abundance profiling. Moreover, though QIIME2 performed well in some prior benchmarking studies [4, 37], Lu et al.[28] demonstrated that Kraken2 [51] is 300 times faster than QIIME2 and achieves higher accuracy therefore we focused on comparing to Kraken2. The software versions for the tools tested in the Results are Cliffy (v2.0.0), Kraken2 (v2.1.2), and Bracken (v2.9).

### Cliff compression reduces index size by orders of magnitude

We indexed the SILVA SSU NR99 database (510,508 rRNA gene sequences, 1.2 GB) to classify 16S rRNA reads and observed a substantial compression ratio (249-275x) for all three index types using different digestion methods (Table 1). For the minimizer index, the average profile storage decreased from *d* = 9118 values to 5.137 pairs in the cliff-compressed profile, leading to a significant reduction in index size. Despite this substantial reduction, Kraken2’s index remained considerably smaller than Cliffy’s index (144 MB vs 4.266 GB).

### Query time on 16S rRNA read datasets

We used both Cliffy and Kraken2 to perform taxonomic read classification over these datasets and measured the running time for both. We observed that, while Cliffy was slower than Kraken2 in all its configurations, configuring Cliffy to use its minimizer-based index yielded a running time that was the closest to Kraken2’s, being on average 4.5x slower than Kraken2 (Figure 2). By contrast, Cliffy’s other modes are closer to 10–11x slower. This result is consistent with previous studies [3, 16] that showed that using minimizers as well as a minimizer-based alphabet can help to speed up read classification.

**Figure 2.**
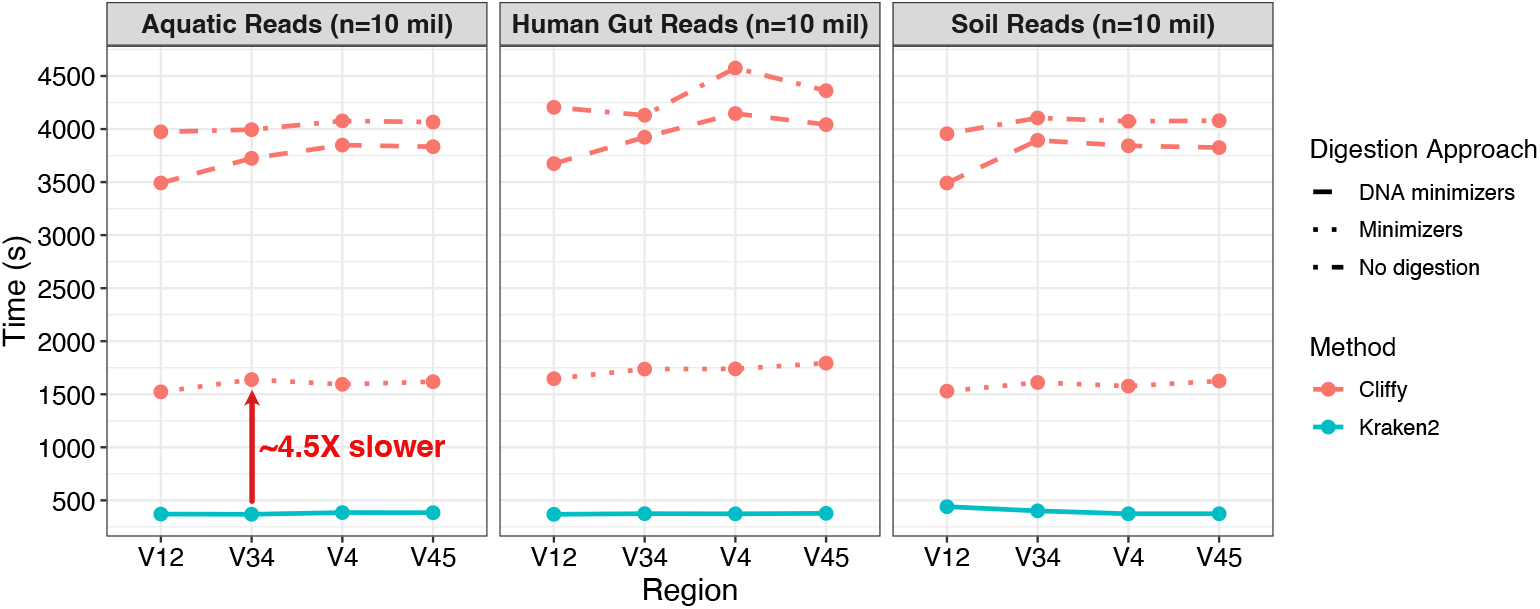
Shows the time required for Cliffy and Kraken2 to classify 10 million simulated Illumina reads (simulated with `art_illumina` [23] using MiSeqv3 error model, 250 bp paired-end). We simulated reads for three different biomes and for each one, simulated reads from four different hypervariable regions.

### Read-level accuracy comparison to Kraken2

Having obtained taxonomic classifications for the simulated reads, we next compared Cliffy’s classification accuracy to Kraken2’s. While we assess accuracy at all taxonomic levels, we are chiefly concerned with accuracy at the lowest level, which for our 16s RNA taxonomy was the genus level.

Measuring accuracy requires special consideration for the case where a tool’s classification is non-specific (i.e. not at a leaf of the taxonomic tree) but also correct, in the sense that read’s genome of origin is below the node reported by the classifier. In particular, we consider four possibilities: <u>True Positives</u> (TPs) are cases where the read is classified to the correct leaf node of the taxonomy. <u>False Positives</u> (FPs) are cases where the read is classified to a node that is neither the correct leaf node, nor an ancestor of the correct leaf node. <u>Vague Positives</u> (VPs) are cases where the read is classified to an ancestor of the correct node. Finally, <u>False Negatives</u> (FNs) are cases where the read is not classified at all by the tool, despite originating from one of the genomes in the taxonomy. Because all simulated reads have their true origin in a node in the taxonomy, True Negatives (TN) are not relevant here. Note that Cliffy reports classifications only at the taxonomic level that is specified by the user; in this case, that was the genus level. Thus, Cliffy does not report any vague positives.

To compute accuracy, we divide TP by the total number of reads in the dataset. This alone is a useful statistic for Cliffy, but for Kraken2 we additionally report a result for accuracy that includes both TPs and VPs in the numerator. Kraken2 results are shown as a range of values, representing a range of accuracy that depends on whether VPs are considered true or false (grey ribbon in Figure 3).

**Figure 3.**
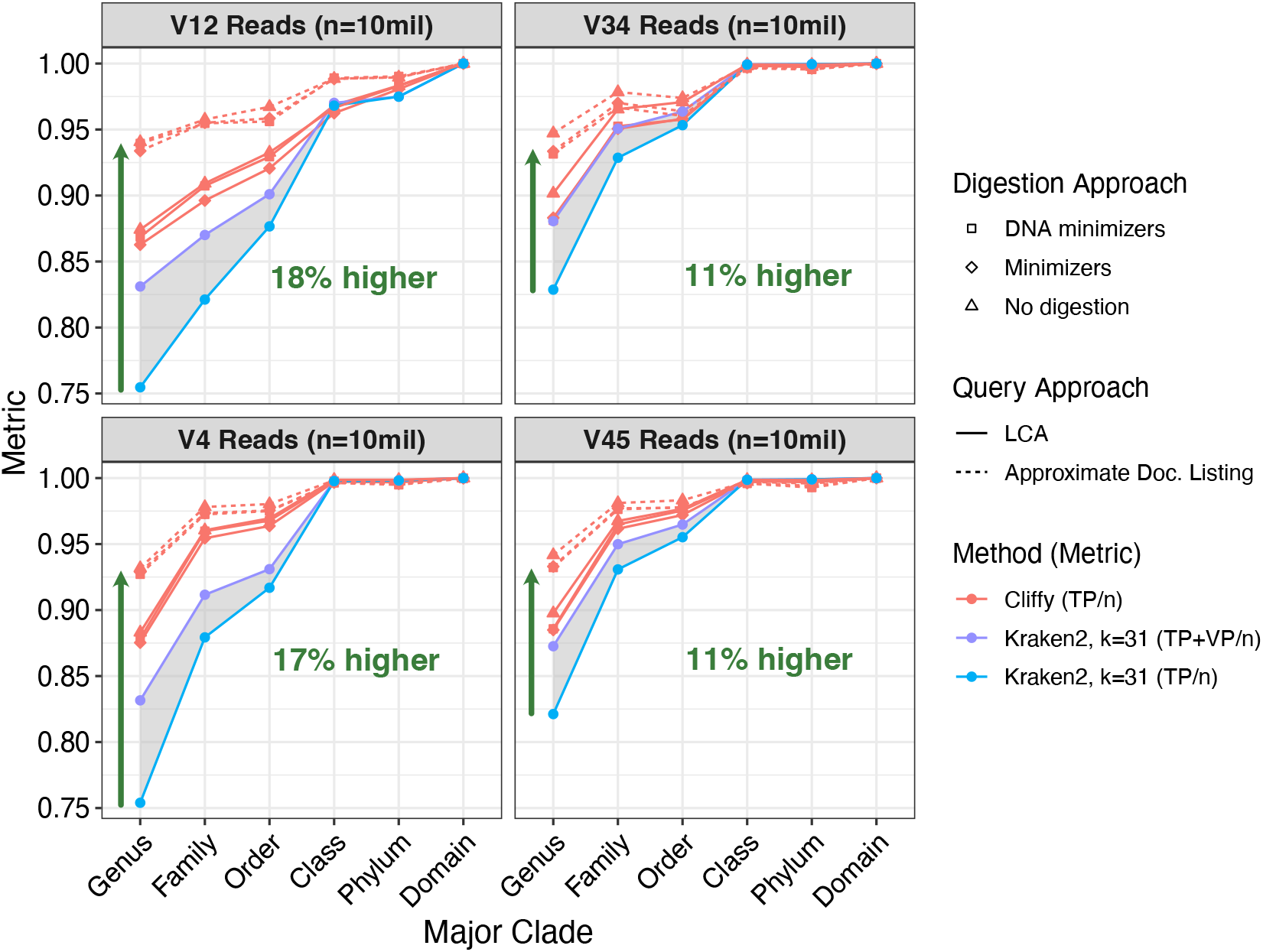
Shows the read-level classification (*n* = 10 million Illumina 250 bp paired-end) accuracy for Cliffy and Kraken2 on the Aquatic dataset using four different hypervariable regions at different levels of the tree. Each sub-plot shows Cliffy’s performance when using three different types of digestion strategy, and two different query approaches: lowest common ancestor and approximate document listing. The grey shaded region ranges from Kraken2’s accuracy when treating all vague positives (VP) as incorrect to treating them all as correct.

Consistently across the four hypervariable regions, Cliffy had higher accuracy than Kraken2. Cliffy’s accuracy percentages ranged from 11 to 18 percent points higher than Kraken2’s at the genus level. This result was consistent across the other two biomes as well (Appendix A.5). We also observed that Cliffy’s more detailed approximate document listing query was consistently more accurate than its lowest common ancestor query. Finally, when comparing the different digestion approaches, we saw that the default minimizer scheme of *k* = 4 and *w* = 11 yielded accuracy results that were near identical to those using the full input text with no digestion.

To better understand why Cliffy had a higher accuracy than Kraken2, we focused on the four datasets for which the accuracy gap was the largest. We visualized where in the taxonomy Kraken2 was identifying the lowest common ancestor nodes for the minimizers it identified from the reads (Figure 4). When comparing reads that Kraken2 classified correctly versus incorrectly (TP vs. FP), we see that Kraken2 identifies less lowest common ancestor nodes at the genus level and instead it finds more at the domain level. Across these four datasets, we see about 30% of the minimizers are not found in the database (i.e. k is too long) and 30-50% of the minimizers are at the domain level (i.e. k is too short). This data suggests the minimizer length used by Kraken2 is not always ideal, whereas Cliffy’s ability to identify variable length matches allows it to overcome this limitation.

**Figure 4.**
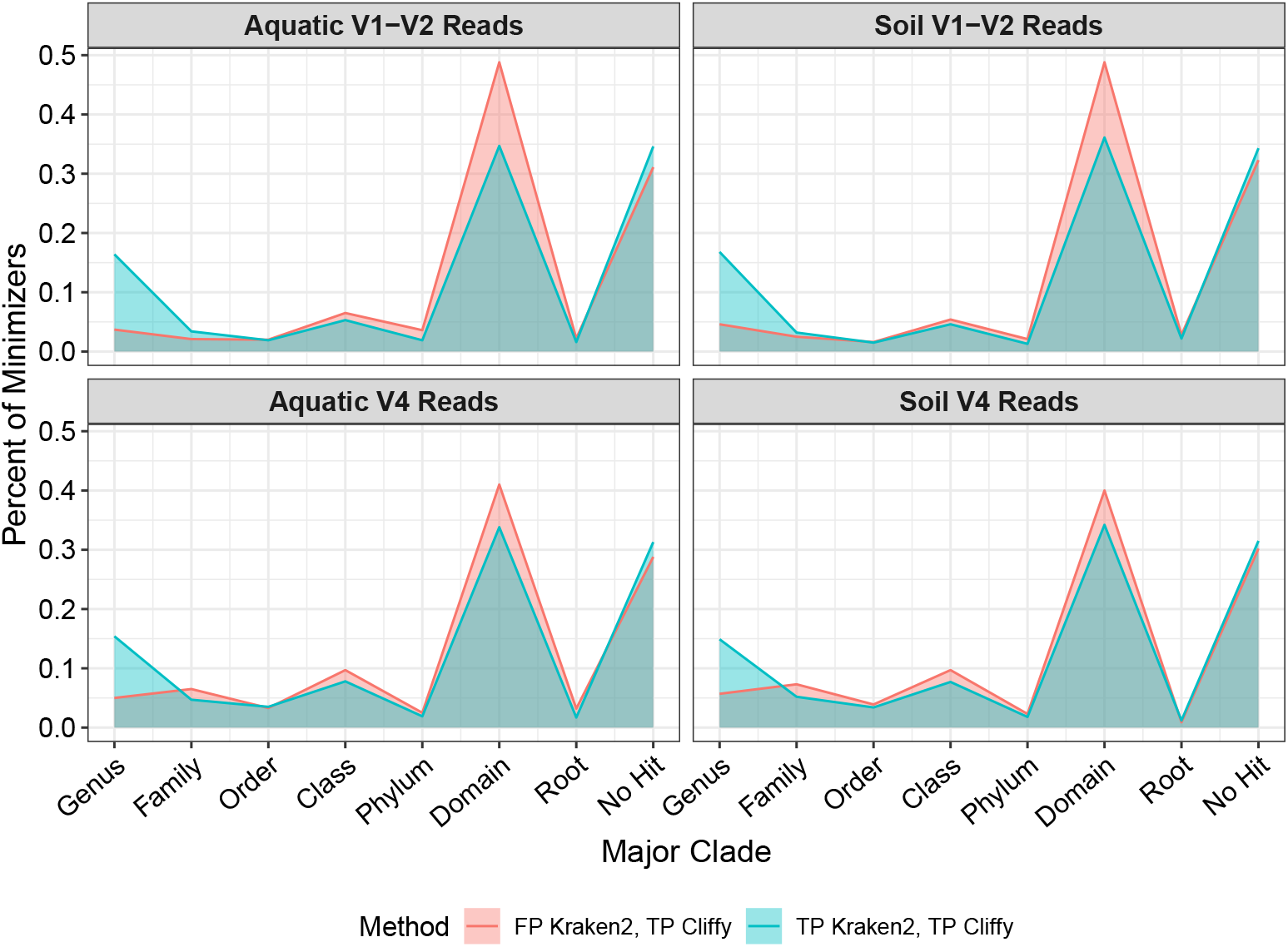
Focuses on the four datasets with the largest accuracy gap between Kraken2 and Cliffy (using LCA query and not approximate document listing). Looking only at the reads that were false positives for Kraken2 and true positives for Cliffy or reads that were true positives for both tools, we added up the number of minimizers that Kraken2 identified at each level of the taxonomy as well as minimizers that did not hit anything in the database to understand why Kraken2’s accuracy was lower than Cliffy despite both tools using the LCA query.

### Genus abundance estimation using Cliffy

The results discussed so far assessed accuracy at the granularity of individual reads. We further compared these tools at the dataset level, comparing their ability to correctly quantify the abundance of each genus. Figure 5 shows the estimated genus-level abundances for the 100 genera simulated in the Aquatic dataset. For all four regions, Cliffy generated a distribution with the lowest Bray-Curtis Distance (BCD) from the true distribution. That trend held as well for the other eight datasets (Appendix A.6).

**Figure 5.**
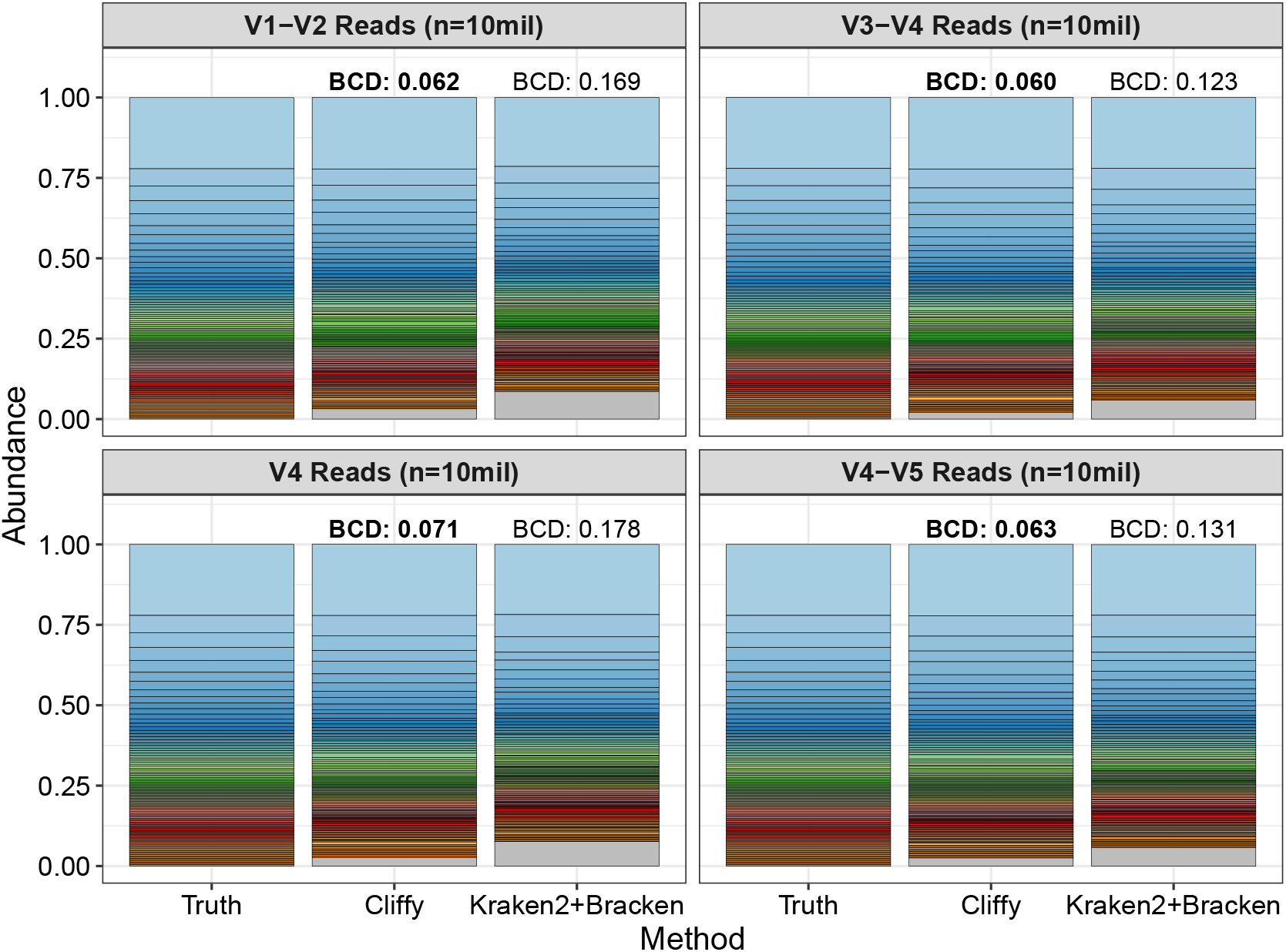
Shows the genera abundance results for Cliffy and Kraken2+Bracken for the Aquatic dataset compared the truth distribution. Each stacked bar is compared quantitatively with the truth distribution through Bray-Curtis Distance (BCD). The method with the closest distribution to the truth (i.e. lowest BCD) is distinguished with a bold BCD value. The grey section of the Cliffy and Kraken2 bars correspond to predicted genera that are not truly present in the dataset.

## 4 Discussion

We introduced Cliffy, which uses a full-text compressed index to perform accurate 16S rRNA gene classifications. We described a novel compression scheme for the document array profiles [1], making it far more practical for taxonomic classification. Its use of full-text indexing, allows Cliffy to find exact matches of variable length, allowing for greater accuracy compared to Kraken2 [51].

The document array profiles [1] was the first document listing data structure to have a complexity (*𝒪* (*rd*)) that depended only on *r* and not *n*, making it particularly practical for repetitive collections like pangenomes. Here we proposed cliff compression, which greatly reduces index size at the expense of enabling only taxonomic classification queries and not full document-listing queries. The compressed index reaches an average size of Θ(*r* log *d*), which we justified using a random model. Further, we showed that we tend to shrink the index somewhat more than expected in practice. This vast reduction in space makes Cliffy far more practical for taxonomic classification than any past full-text indexing approach.

*k*-mer based tools like Kraken2 work by pre-selecting a particular value for *k* (e.g. *k* = 31), but there is no reason to expect any particular value of *k* is universally appropriate for all sequencing technologies and all portions of the tree of life. Here we saw evidence both of scenarios where a fixed value of *k* was too long (30% minimizers hit nothing in the database), and where it was too short (30%-50% of minimizers have an LCA at the domain level). In the future, it will be important to continue studying full-text indexes with no pre-selection of *k*, as this could be critical to enabling high accuracy over time and over various technologies and clades.

Despite the query optimizations and compression, the main weakness of Cliffy is the lower computational efficiency compared to the *k*-mer based Kraken2 method. Currently, Kraken2 is about 4.5x faster classifying reads compared to Cliffy when using minimizer digestion. Also, Kraken2’s index is about 30x smaller (4.266 GB vs 144 MB). While this work represents a major closing of the index-size gap, we will continue to look for opportunities to close the gap in terms of both speed and index size. To reduce query time, we could consider using faster rank queries that could potentially speed up the backward search step in Cliffy’s code for finding pattern matches [9, 14]. Further speed improvement could come from using an alternative data structure for the run-length-compressed BWT, such as the move-structure [36, 53] that allows for faster backward search (up to 30X faster than current approaches) at the cost of a larger index.

To reduce Cliffy’s index size, we can exploit the inherent feature of cliff-compression where the stored profile values (*CP*_*𝓁r*,*r𝓁*_[*i*][*j*]) are strictly increasing, therefore, we could use a delta-encoding scheme to reduce the memory needed to store them. Recent works in k-mer indexes have used exploited the repetitiveness of colors to reduce the size of their indexes [17]. It is possible a similar approach can be applied to the cliff-compressed document array profiles if groups of documents ids tend to be stored together after applying the compression.

## Acknowledgements

This work was performed at the Advanced Research Computing at Hopkins (ARCH) core facility (https://www.arch.jhu.edu/), which is supported by the National Science Foundation (NSF) grant number OAC 1920103. This work was also supported by the NSF grant number DBI-2029552 and by the National Institutes of Health grant numbers R01HG011392, R35GM139602, and T32GM119998.

## 5 Software Availability

The Cliffy software tool can be obtained via GitHub at this link: https://github.com/oma219/cliffy. The experiment code used to generate the results for this paper can be found at this link: https://github.com/oma219/cliffy-experiments.

## 6 Author Contributions

OA and BL conceived of the cliff compression idea. CB did foundational work on the *r* index and and document array listing. OA developed the Cliffy software and ran the experiments. OA wrote and all authors edited the manuscript. All authors approved the manuscript.

## 7 Conflicts of Interest

The authors declare no conflicts of interest.

## A Appendix

### A.1 Proof of equivalence to harmonic number

Equation 2 stated this relationship between our recursive formula for **E**[*S*_*n*_] and the harmonic number.

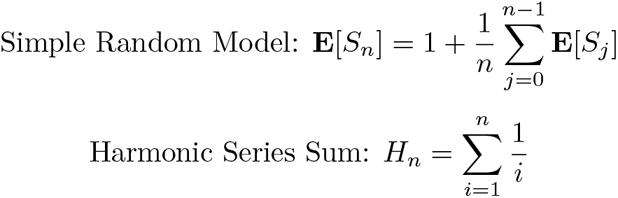

We now show that these are equivalent using induction.

**Proof**.

**Base case** Letting *n* = 1 for both equations:

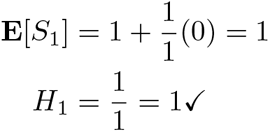

**Induction step**. We now assume **E**[*S*_*n*_] = *H*_*n*_ and show that **E**[*S*_*n*+1_] = *H*_*n*+1_.

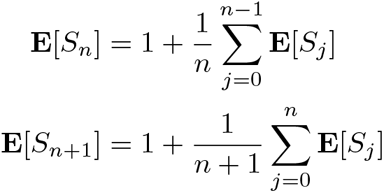

Now we rewrite **E**[*S*_*n+*1_] in terms of **E**[*S*_*n*_]:

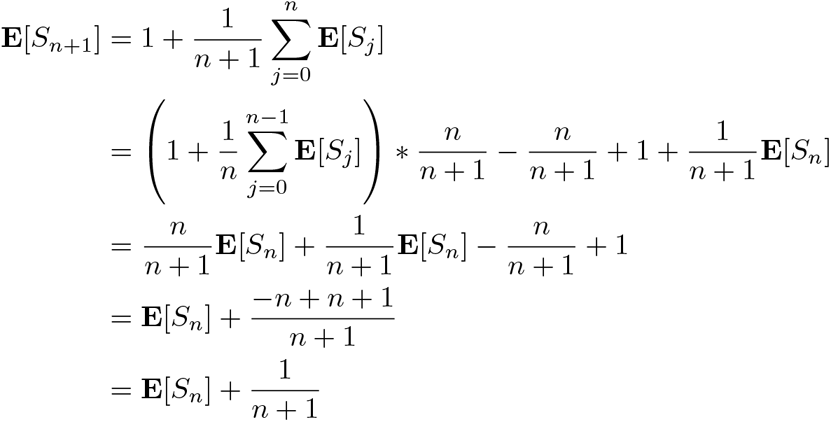

Now apply the assumption of *S*_*n*_ = *H*_*n*_.

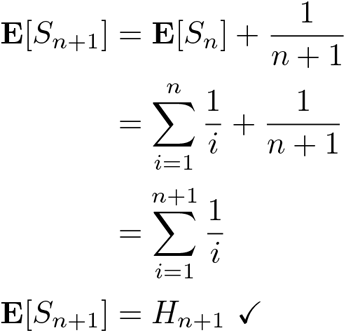

### A.2 Proof of membership in Θ(log *d*)

In Section A.1, we proved the expectation of our recurrence sum for a simple random model is equivalent to the harmonic series sum (**E**[*S*_*n*_] = *H*_*n*_). Next, we want to show that *H*_*n*_*∈* Θ(log *n*) in order to prove the average space complexity for the cliff-compressed document array profiles.

**Proof**. Using Stolz–Cesàro’s rule (L’Hôpital’s rule for sequences), we can show that *H*_*n*_ ∈ Θ(log *n*) by proving that as *n → ∞* the limit of the ratio of the functions is equivalent to a constant. If so, that means we can identify constants *c*_1_, *c*_2_ such that *c*_1_ log *n ≤ H*_*n*_ *≤ c*_2_ log *n* thereby proving that *H*_*n*_ *∈* Θ(log *n*).

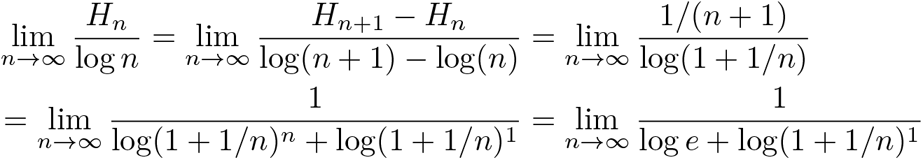

Here, we know that as *n* approaches ∞ the expression log(1 + 1*/n*) will approach log(1) which is equivalent to 0.

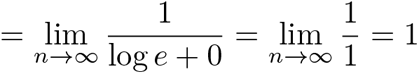

Therefore, we have shown that *H*_*n*_ *∈* Θ(log *n*), and based off of the result from the previous section (A.1), we have also shown that **E**[*S*_*n*_] *∈* Θ(log *n*)

### A.3 Running Kraken2 and Braken

Here are command used to build both the Kraken2 and Bracken indexes for the SILVA database.

~~~
kraken2-build --db silva --special silva --threads 1
bracken-build -d silva -t 1 -l 250 -k 35
~~~

Here are the commands used to run the Kraken2 query followed by using Bracken to do genus abundance estimation.

~~~
kraken2 --db silva
        --threads 1
        --report output.kreport2
        --paired mate_1.fastq mate_1.fastq
        > output.kraken2
bracken -d silva
        -r 250
        -l G
        -i output.kreport2
        -o output.bracken
~~~

### A.4 MicrobeMixer: Simulating realistic 16S rRNA reads

The code to use `MicrobeMixer` can be found at https://github.com/oma219/MicrobeMixer. There are two main steps; the first step is to summarize public data to identify the top 100 genera for a particular biome and the second step is to simulate reads for that biome using `art_illumina` [23].

Here is an example of how to run `MicrobeMixer` to generate an abundance file that contains the top 100 genera found in human gut.

~~~
python3 sim_16s_reads.py stats
                   --biome root:Host-associated:Human:Digestive system
                   --taxonomy ../data/tax_slv_ssu_138.1.txt
                   --output abundance.tsv
~~~

The previous code generates a file called abundance.tsv. Next we will use that file to generate simulate reads from those genera.

~~~
python3 sim_16s_reads.py simulate
                   --biome-abundance abundance.tsv
                   --silva-ref ../data/SILVA_138.1_SSURef_NR99_tax_silva.fasta
                   --silva-taxonomy ../data/tax_slv_ssu_138.1.txt
                   --primers ../data/V3_V4_primers.txt
                   --temp-dir <OUTPUT_DIR>
~~~

This command will generate two FASTQ files and file called seqtax.txt that contains the mapping for each genus id (1 to 100) to the taxonomic path for that genus. Here is the command that `MicrobeMixer` uses to simulate reads using `art_illumina`.

~~~
art_illumina -ss MSv1 -amp -p -na -i genus_{genus_num}_seqs.fna -l 250
             -f {coverage} -o genus_{genus_num}_reads_
~~~

Additionally, the primers used by `MicrobeMixer` to extract the hypervariable regions are list below which were same ones used by Almeida et al [4]:

~~~
V1-V2 F: AGMGTTYGATYMTGGCTCAG        V4-V4 F: GTGCCAGCMGCCGCGGTAA
V1-V2 R: GCTGCCTCCCGTAGGAGT          V4-V4 R: GACTACHVGGGTATCTAATCC
V3-V4 F: CCTACGGGNGGCWGCAG           V4-V5 F: GTGCCAGCMGCCGCGGTAA
V3-V4 R: GACTACHVGGGTATCTAATCC       V4-V5 R: CCCGTCAATTCMTTTRAGT
~~~

### A.5 Read-level classification results for additional datasets

Figures 6 and 7 show the read-level classification on the Human Gut and Soil datasets.

**Figure 6.**
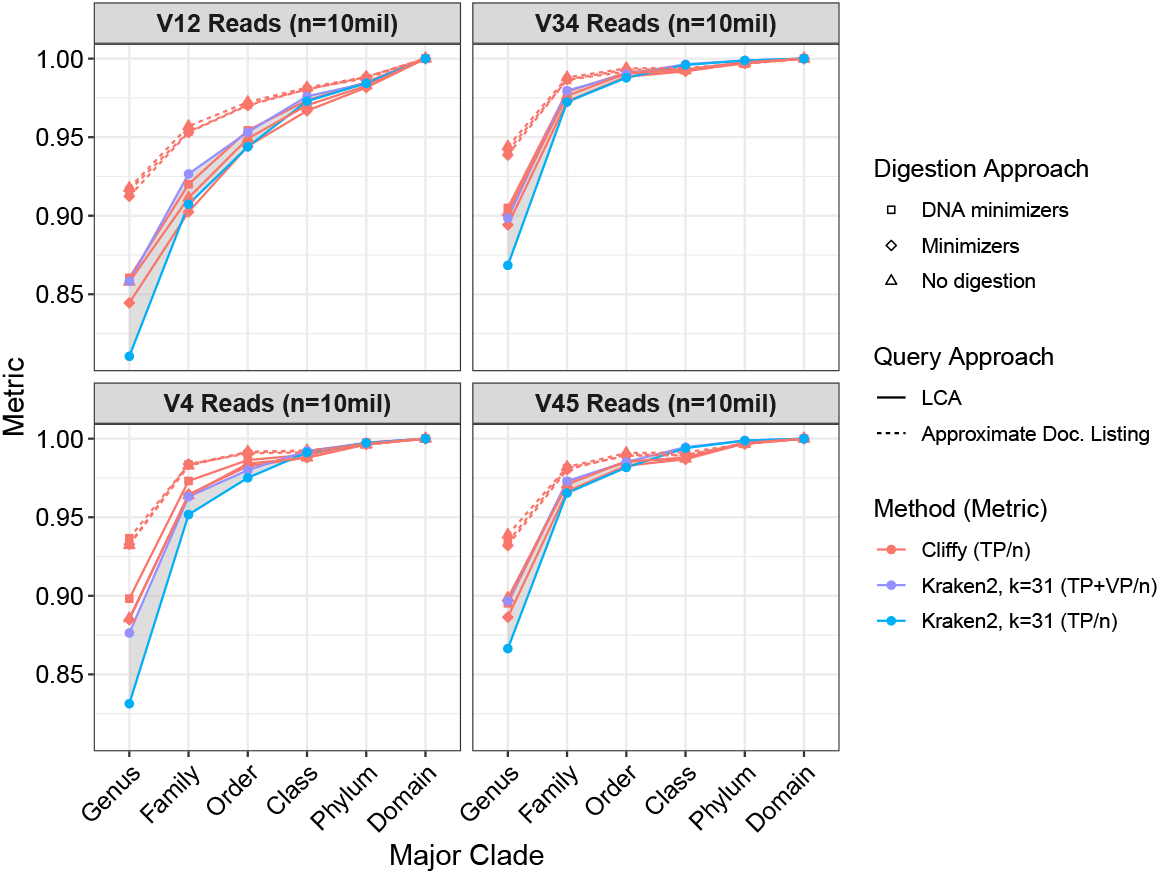
Read-level classification results across different levels of taxonomy on the Human Gut datasets corresponding to four different hypervariable regions. Each dataset consisted of 10 million 250-bp paired-end Illumina reads.

**Figure 7.**
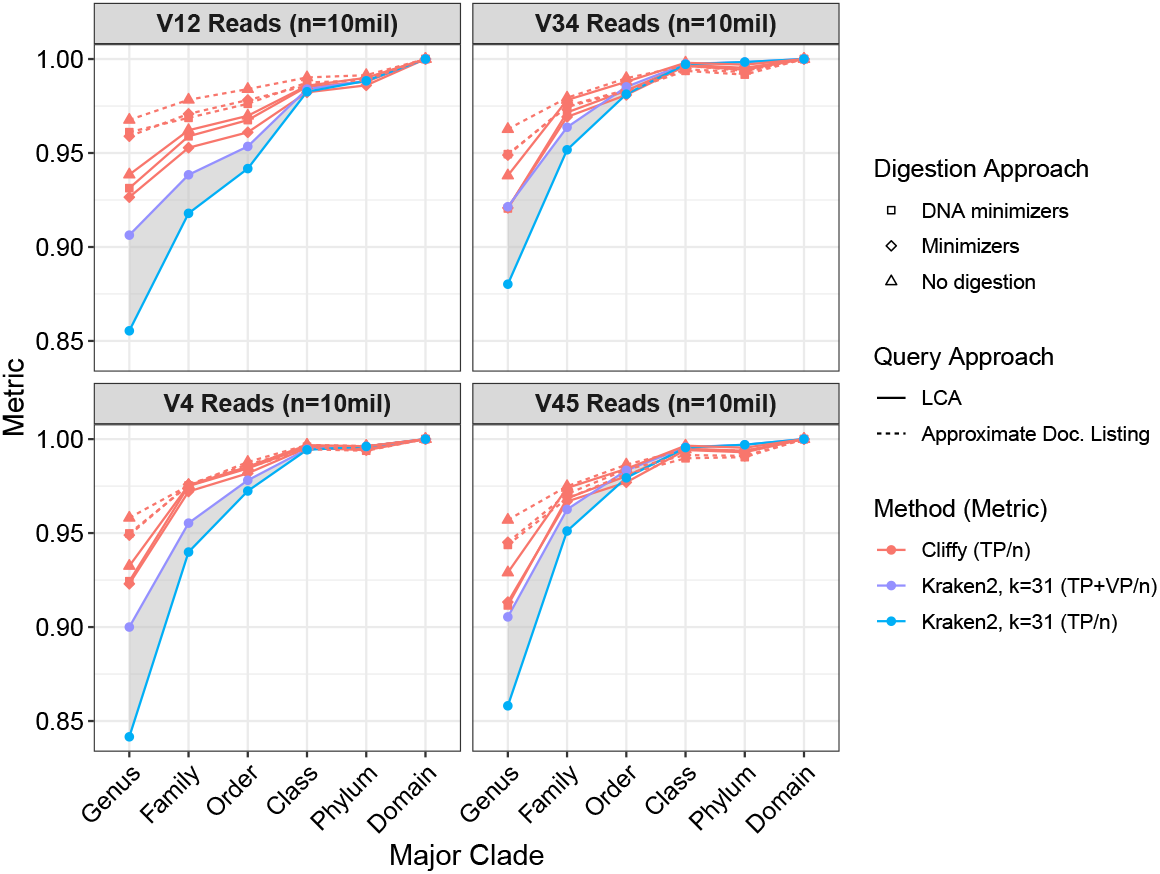
Read-level classification results across different levels of taxonomy on the Soil datasets corresponding to four different hypervariable regions. Each dataset consisted of 10 million 250-bp paired-end Illumina reads.

### A.6 Genera abundance results for additional datasets

Figures 8 and 9 show the genera abundance estimation and Bray-Curtis distances on the Human Gut and Soil datasets.

**Figure 8.**
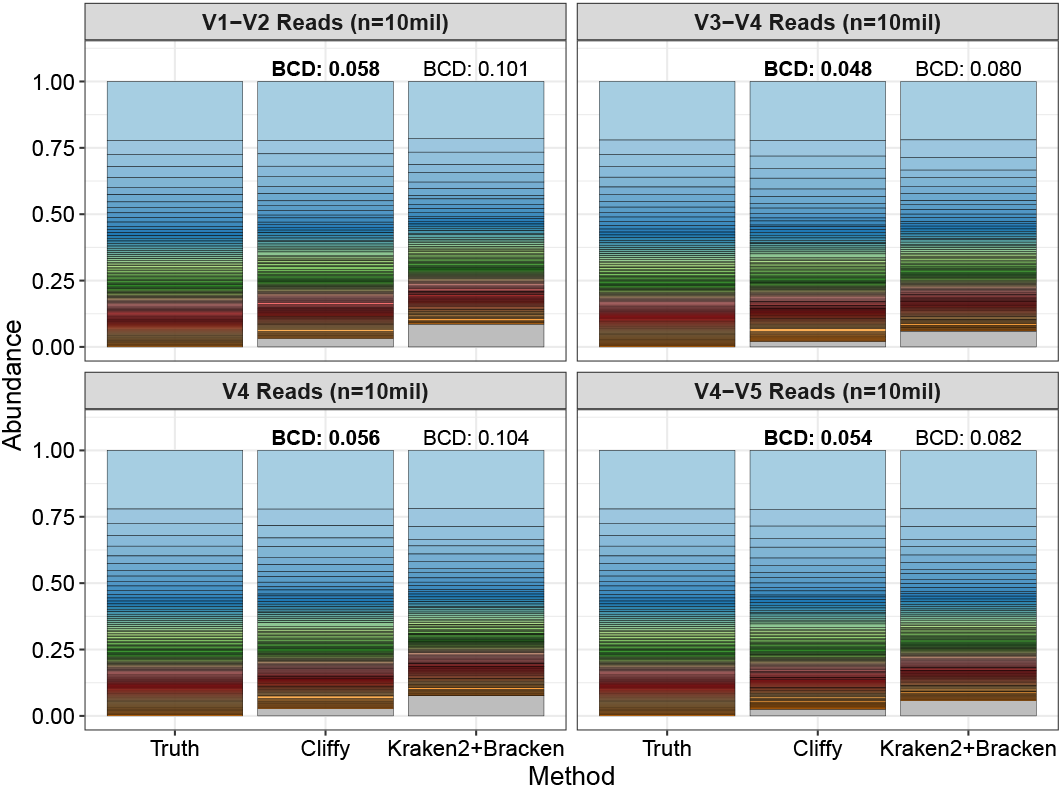
Genera abundance estimation for the 100 genera present in each of the simulated datasets for the Human Gut microbiome. The grey bar present in the Cliffy and Kraken2 bars present the percentage of reads classified to genera not present in the true set. Each bar is annotated with the Bray-Curtis distance to the true distribution (lower means closer).

**Figure 9.**
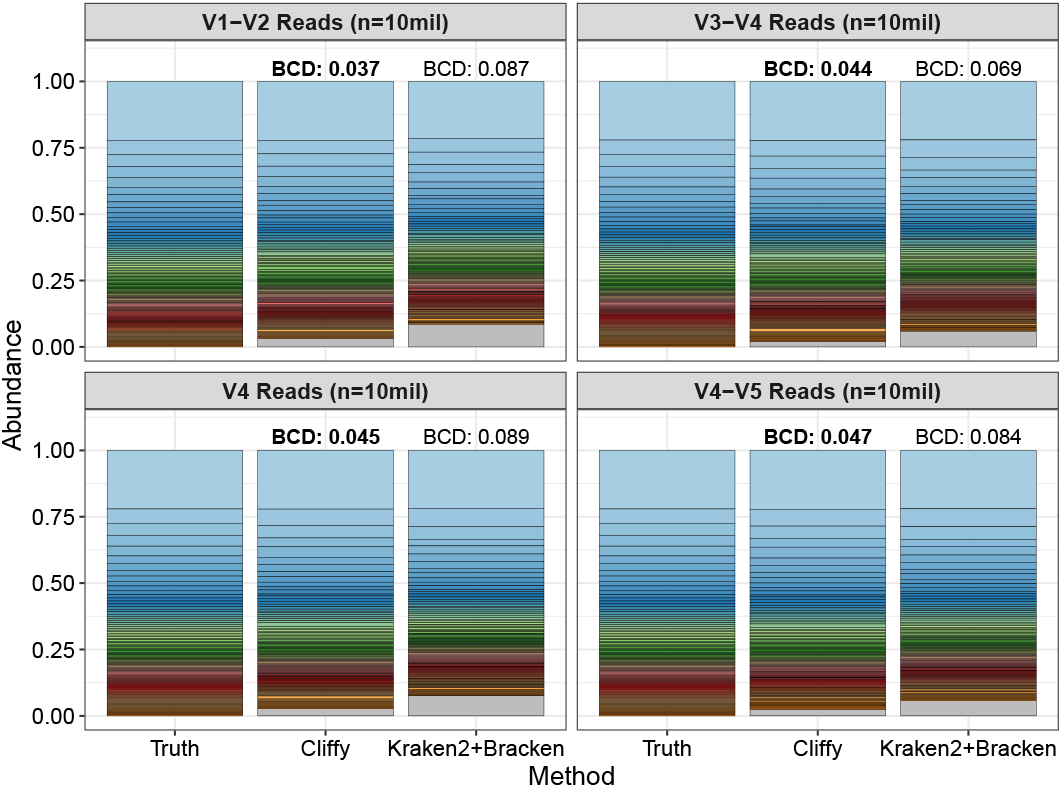
Genera abundance estimation for the 100 genera present in each of the simulated datasets for the Soil microbiome. The grey bar present in the Cliffy and Kraken2 bars present the percentage of reads classified to genera not present in the true set. Each bar is annotated with the Bray-Curtis distance to the true distribution (lower means closer).

## Notes

https://github.com/oma219/cliffy

https://github.com/oma219/cliffy-experiments

### Competing Interest Statement

The authors have declared no competing interest.

